# Localization of Albumin with Correlative Super Resolution Light- and Electron Microscopy in the Kidney

**DOI:** 10.1101/2024.09.16.613238

**Authors:** Alexandra N. Birtasu, Utz H. Ermel, Johanna V. Rahm, Anja Seybert, Benjamin Flottmann, Mike Heilemann, Florian Grahammer, Achilleas S. Frangakis

## Abstract

The functioning of vertebrate life relies on renal filtration of surplus fluid and elimination of low-molecular-weight waste products, while keeping serum proteins in the blood. In disease, however, there is leak of serum proteins and tracing them to identify the leaking position within tissue with a nanometer resolution poses a significant challenge. Correlative microscopy integrates the specificity of fluorescent protein labeling into high-resolution electron micrographs. Using chemical tagging of albumin with synthetic fluorophores we achieve protein-specific labeling that preserve their post-embedding fluorescence after high-pressure freezing and freeze-substitution of murine kidney tissue. Using advanced registration techniques for super-resolution correlative light and electron microscopy, we can localize the labeled albumin with a high precision in the x-y plane of electron micrographs and cartograph its distribution. Thereby we can quantify the albumin concentration and measure a linear reduction gradient across the kidney filtration barrier. Our study shows the feasibility of combining different microscopy contrasts for tracing fluorescently labeled protein markers with super resolution in various tissue samples and opens new perspectives for correlative imaging in volume electron microscopy.

## Introduction

In vertebrates, the kidneys play a crucial role in maintaining precise water and electrolyte balance within the body. They efficiently remove low-molecular-weight waste products from significant amounts of plasma while simultaneously preserving essential proteins within the bloodstream. This filtration process primarily occurs within microvascular units known as glomeruli, which comprise three distinct layers: a monolayer of fenestrated endothelial cells in the capillary space, a robust glomerular basement membrane (GBM), and specialized epithelial cells the podocytes in the urinary space (Farquhar et al., 1961; Yamada, 1955). The podocytes have a unique intercellular junction, the slit diaphragm (SD), which connects to adjacent podocytes [3]. Numerous kidney diseases are linked to dysfunction of the SD, with pathological alterations in SD evident in acquired and hereditary conditions associated with proteinuria. Such conditions include minimal change disease, congenital nephrotic syndrome, nephrotic syndrome, and diabetes [4,5]. This underscores the critical need to comprehensively understand the kidney filtration system at the molecular level, paving the way for the development of effective therapies targeting proteinuria.

The SD has been the subject of extensive study utilizing various techniques. Advanced light microscopy coupled with electron microscopy tracers has enabled the investigation of the filter’s selectivity properties under physiological conditions by visualizing protein distribution at the filtration barrier [6]. Elaborate physical models have attributed the size and charge selectivity of the glomerular filtration barrier to streaming potentials, which effectively prevent most of the filtrate from entering the filtration barrier at the endothelial cells [7]. Lately, the cryo-electron tomography (cryoET) of vitreous lamellae allowed for the visualization of the SD at native conditions [8]. The cryoET structure highlights the significance of the SD in its role in providing mechanical resistance against blood pressure, but also providing dedicated holes (previously called “pores”) highlighting its multifaceted functionality [8].

Previously we established a correlative super resolution Light- and Electron Microscopy (CLEM) protocol for localizing proteins in cultured cells [9]. Here we extend this protocol for tissue samples. We apply it in murine kidney tissue and measure the precise location of fluorescently labelled albumin at the glomerular filtration barrier. We estimate the distribution of albumin with respect to the individual compartments and estimate its distribution. This feasibility study shows that different protein pathways can be traced by this technique and makes an estimate of the albumin distribution at the glomerular filtration barrier.

## Material and Methods

The research presented in herein complies with all relevant EU, national and regional laws and regulations. We also comply with EU, national and international ethics-related rules and professional codes of conduct.

### Animals for CLEM albumin localization

All experiments were conducted in accordance with the German animal welfare guidelines and the NIH Guide for the Care and Use of Laboratory Animals and were approved by the responsible authority (G-07/13 Regierungspräsidium Freiburg, Germany). C57Bl/6 mice were housed in a specific-pathogen-free (SPF) facility with free access to chow and water and a 12 h day/night cycle.

### CLEM albumin sample preparation – tissue processing

Animals were intraperitoneally injected with ketamine/xylazine for sedation (7 µL/kgBW, 7 g/kgBW). Kidneys were injected intravenously through the retroorbital plexus with 500 µg bovine serum albumin (BSA) (Alexa Fluor™ 555 conjugate, Thermo Fisher Scientific Inc., Waltham, MA, USA). After 60 s, kidneys were dissected together with the abdominal aorta. The empirically derived time interval was used to ensure thorough mixing in the blood, while keeping it short to prevent albumin from being excreted from the kidney. The tissue was cut into small pieces and placed in gold-plated copper specimen carriers with 0.15 μm recess (type 665; Wohlwend, Sennwald, Switzerland) filled with 20% dextran. The tissue sections were high-pressure frozen in an HPM-010 (Abra Fluid, Widnau, Switzerland). Freeze substitution was performed in an automated system (EM AFS2, Leica Microsystems, Wetzlar, Germany) with a solution of 2% uranyl acetate (Serva, Heidelberg, Germany), 8% methanol and 2% water dissolved in glass-distilled acetone (Electron Microscopy Sciences, Hatfield, USA) at –90°C for 64 h and at –45°C for 24 h, followed by three washes in acetone at –45°C for 10 min. Tissue sections were sequentially infiltrated with increasing concentrations (10%, 25%, 50% and 75%) of Lowicryl HM20 (Polysciences, Hirschberg an der Bergstrasse, Germany) for 4 h each, while raising the temperature to –25°C in increments of 5°C. Lowicryl was exchanged two times after 12 h and a third time after 48 h. The tissue sections were then UV polymerized for 48 h at – 25°C, followed by another 9 h at 20°C. The tissue sections were observed in a confocal microscope (LSM700, Carl Zeiss, Jena, Germany) to identify fluorescent glomeruli. Distances of glomeruli to tissue edges were measured. Ultrathin sections (40 nm) were cut with an ultramicrotome (Ultracut UC7, Leica Microsystems) and transferred to formvar-coated copper finder grids (Plano).

### Super-resolution imaging

Measurements were conducted on a home-built microscope for single-molecule localization microscopy. In brief, an inverted microscope (IX81, Olympus, Tokyo, Japan) was equipped with a 150× UApo TIRFM oil objective lens (NA 1.45, Olympus) and operated in total internal reflections fluorescence (TIRF) mode. The fluorescence signal was collected on an Andor Ixon Ultra EMCCD chip (DU-897U-CS0-#BV; Andor Technology Ltd, Belfast, Northern Ireland). The average localization precision was 47 nm, which is approximately the width of the SD. While the resolution is influenced by the background noise, it remains better then diffraction limited wide-field microscopy. Samples were imaged in PBS (Polyscience) with 50 – 200 mM MEA (Sigma-Aldrich, St. Louis, Missouri, US, #M6500-25G) added to adjust single-molecule photoswitching of the fluorophore [10]. Alexa555 BSA Albumin (Thermo Fisher Scientific Inc.) was excited with a 561 nm diode laser (Cube; Coherent, Newton, CT, USA). Illumination intensities were adjusted to 0.65 kW/cm^2^ for direct stochastic optical reconstruction microscopy (dSTORM) imaging. We collected 3000 frames with an integration time of 200 ms, an image size of 512×512 px^2^ with a pixel size of 108 nm, and an EM gain of 200.

### Localization and clustering of dSTORM movies

The dSTORM movies were processed using Picasso [11] (version 0.4.11) to obtain single-molecule localizations. In a first step, localizations were found by using the integrated Gaussian maximum likelihood estimation algorithm of Picasso Localize (parameters: baseline = 178 photons, sensitivity = 15.5 photons, quantum efficiency = 0.97, box side length = 7 px, min net gradient of 800–2200 photons). Localizations were drift corrected in Picasso render using redundant cross-correlation with a segmentation value of 500 frames. To remove signal from out-of-focus planes, localizations were filtered by their standard deviation (up to a magnitude of 324 nm) and by their localization precision (up to a magnitude of 108 nm). Localizations were linked with a radius of 74.3 nm, which is four times the localization precision of nearest neighbor-based analysis (NeNA) [12] and with a maximum dark time of 400 ms [13]. Localizations were clustered with the density-based spatial clustering of applications with noise (DBSCAN) algorithm [14]. The radius was set to 74.3 nm and the minimum density to 7 localizations.

### Correlative transmission electron microscopy

Following dSTORM imaging, montaged transmission electron microscopy (TEM) images of the grids were acquired using a 300 kV TEM (Tecnai F30 G2; Thermo Fisher Scientific Inc.) equipped with a charge-coupled device (CCD) camera (UltraScan 4000; Gatan Inc., Pleasanton, CA, USA). Image montages were acquired automatically using SerialEM 3.6.1 [15] at a nominal defocus of –5 µm and a total dose of 5.1 e^-^ Å^-2^. The calibrated pixel size was 10.22 Å (×12,000 magnification).

The montaged TEM images were pre-processed by custom MATLAB scripts (MATLAB 2021b, 2021, The MathWorks, Natick, MA, USA). To generate single overview TEM images, we blended montages using the software package IMOD v4.11.7 [16]. The registration between light and electron microscopy data was performed by selecting control points on both WF and TEM images and computing an affine spatial transformation using MATLAB (MATLAB 2021b, 2021, The MathWorks). Accuracy of the image registration was compared by overlaying mask and images, respectively. Due to distortions of the sample surface by the preparation, we omitted from our analysis any areas of the sample that showed a higher degree of warping (Supplementary Figure 1).

### Binary mask generation and cluster quantification

The luminal capillary, luminal urinary space, and glomerular filtration barrier were segmented in the EM images using Fiji to create binary masks [17]. The masks were transformed to fit the rendered image of the clusters, as described above. Cluster densities inside the three segmentations were determined by counting the number of cluster centers inside the respective segmented areas and dividing the counts by the area of the segmented mask with a custom script. The mean and standard deviations were calculated. Paired t-tests were applied to compare the means of the 28 segments in four field of views. All distributions were significantly drawn from a normally distributed population, as determined by a Kolmogorov-Smirnov test (*α* = 0.05). Levels of significance were classified as p > 0.05, no significant difference (n.s.); p < 0.05, significant difference (*); p < 0.01, very significant difference (**); and p < 0.001, highly significant difference (***).

## Results

### Preparation of the animals for CLEM

CLEM allows functional assignment of proteins prior to their EM analysis. Here, we analysed murine kidney tissue (microbiopsies (n=5)), injected with bovine serum albumin labelled with Alexa Fluor™555, which was high-pressure frozen, and freeze substituted (Fig. 1). Ultrathin sections (40 nm) were subjected to direct Stochastical Optical Reconstruction Microscopy (dSTORM) data acquisition for the investigation of albumin (approximately 66 kDa) distribution and relocated regions were analysed by TEM imaging recording large landscapes of the glomerular section. The post-embedding fluorescence was well preserved, and the subcellular structural preservation was appropriate allowing for sufficient contrast for a precise image registration owing to the rapid tissue preparation protocol.

**Fig. 1:**
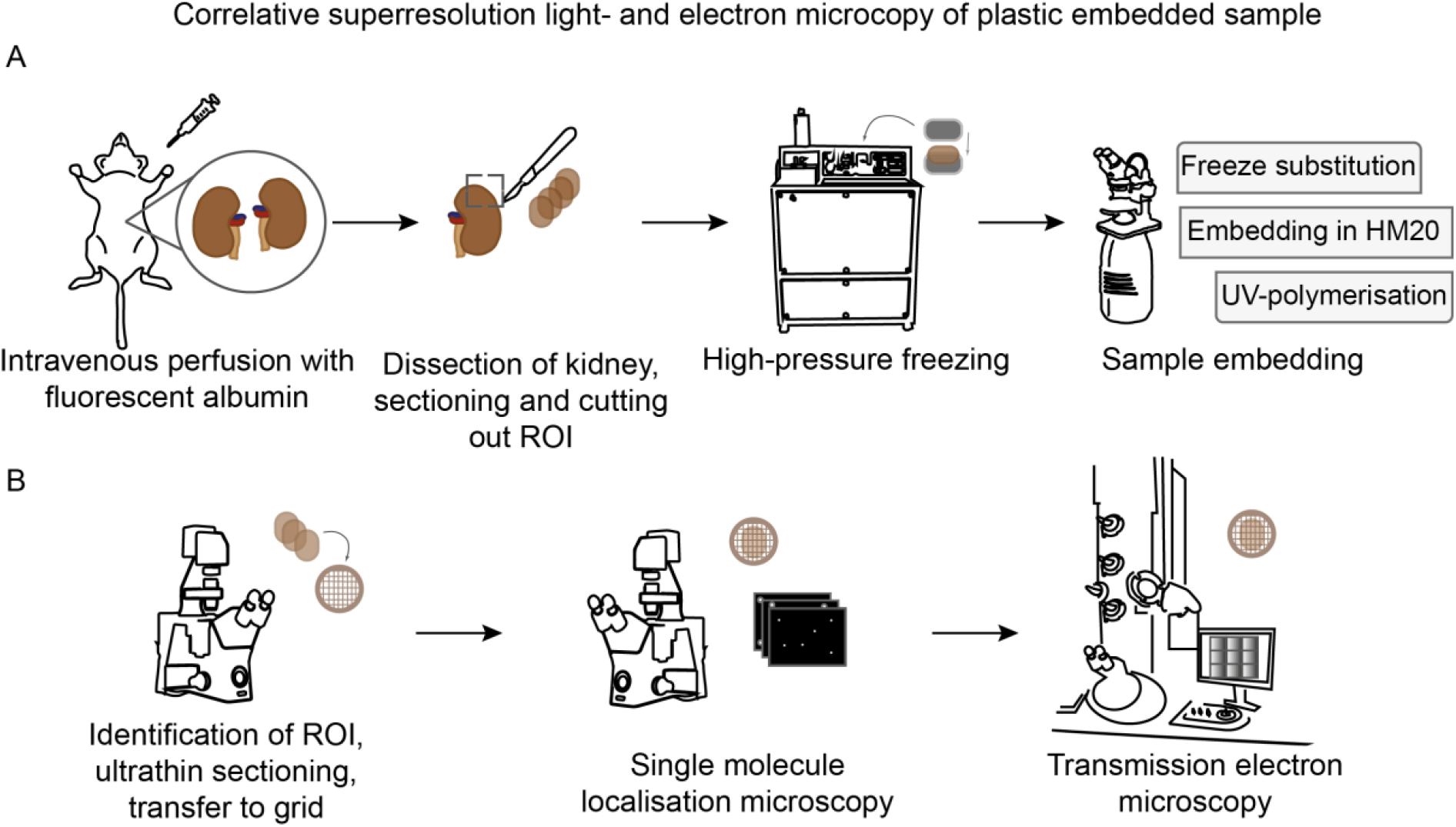
Preparation and imaging pipeline for plastic-embedded kidney tissue. (A) Fresh mouse kidneys, that were injected with fluorescently labelled bovine serum albumin; were dissected and sections of the cortex (ROI) were used for high-pressure freezing, followed by freeze substitution and plastic embedding. (B) Glomeruli representing the regions of interest (ROI) were identified by light microscopy imaging. After thin sectioning and transfer onto electron microscopy (EM) grids, the samples are imaged by direct stochastic optical reconstruction microscopy (dSTORM), followed by transmission electron microscopy.

### Image registration of the different microscopy contrasts

The precision of superposition of the microscopy images is important for accurately localizing the labeled proteins within the EM micrographs. For this, we used the capillary boundaries of the tissue sections that were sufficiently contoured in the widefield (WF) images that accompanied the super resolution images. The indirect contouring of the capillaries is possible since the fluorescent dye does not diffuse into the tissue. Therefore, the non-fluorescent regions served as boundaries, essentially employing a negative selection approach. Those were assigned to features from the structurally preserved glomerular regions in TEM micrographs. The affine transformations were applied to to generate an overlay of dSTORM and EM images (Fig. 2) necessary to compensate for distortions due to the different orientation of the sample in the optical path as well as sample shrinkage due to electron irradiation. Density-based spatial clustering of applications with noise (DBSCAN) analysis was performed to identify albumin clusters.

**Fig. 2:**
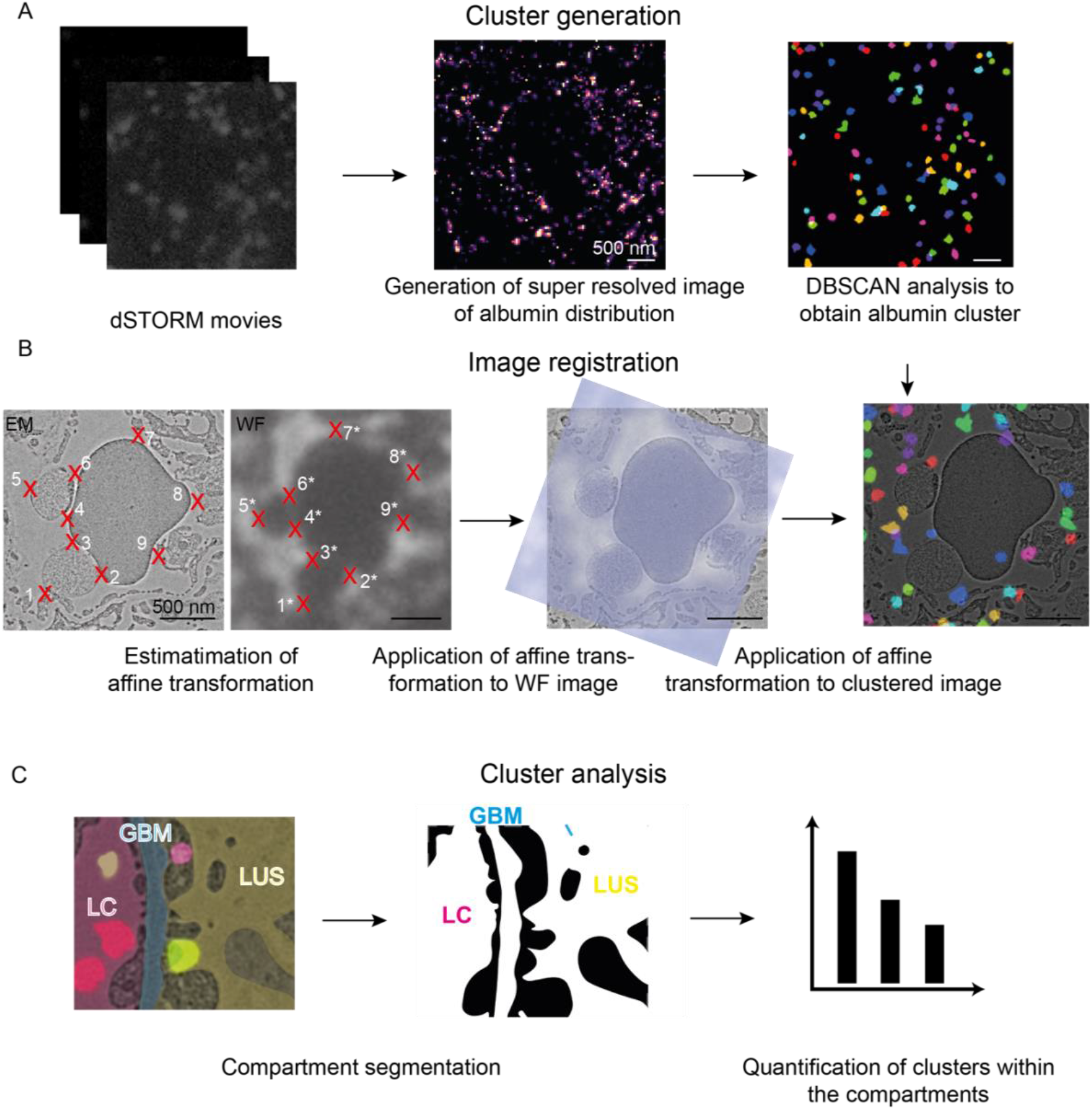
Image analysis. (A) Cluster registration: The dSTORM movies were processed to obtain single-molecule localizations. Localizations were found, drift corrected, filtered by their standard deviation and rendered to generate the super-resolved image. Albumin localizations were clustered with the DBSCAN algorithm. (B) Image registration: Micrographs were preprocessed (bandpass filtered and normalized for compensating different intensities). Subsequently positions with high contrast (control points) were identified in both the EM and the WF images (indicated by the red crosses). The registration between light and electron microscopy data was performed by superimposing control points on both widefield and TEM images and computing an affine spatial transformation. This transformation is also applied to the cluster analysis. (C) Cluster analysis: In a last step, binary masks of the different compartments – the luminal capillaries (LC), glomerular basement membrane (GBM) and lumen of the urinary space (LUS) – are generated and used for cluster quantification.

### Visualisation of albumin at the SD

Next, we analysed the static filtration properties of the SD by CLEM under capillary no-flow conditions. TEM images revealed large landscapes of the glomerular sections, showing all characteristic landmarks, namely the lumen of glomerular capillaries (LC), the lumen of the urinary space (LUS) and glomerular basement membrane (GBM) (Fig. 3a). Each albumin molecule is potentially labelled by multiple fluorophores, and several albumin molecules can also be in close proximity. To account for these factors and to exclude contaminant blinking events, we grouped the localizations into individual clusters using the DBSCAN algorithm (see Material and Methods). Thus, each cluster may include several albumin molecules that were in proximity, inevitably leading to larger cluster sizes; however, their validity remains correct. Each cluster was assigned to an individual compartment of the glomerulus, respectively (Fig. 3b). The localization of albumin clusters within either the capillaries or the lumen does not correlate with any particular landmark, as the albumin is freely moving within the capillary. The albumin clusters can be seen overall at the glomerular filtration barrier, in particular in between the glomerular endothelial cells (GEC), at the GBM, and between the podocyte (P) foot processes (fp) (Fig. 3b).

**Fig. 3:**
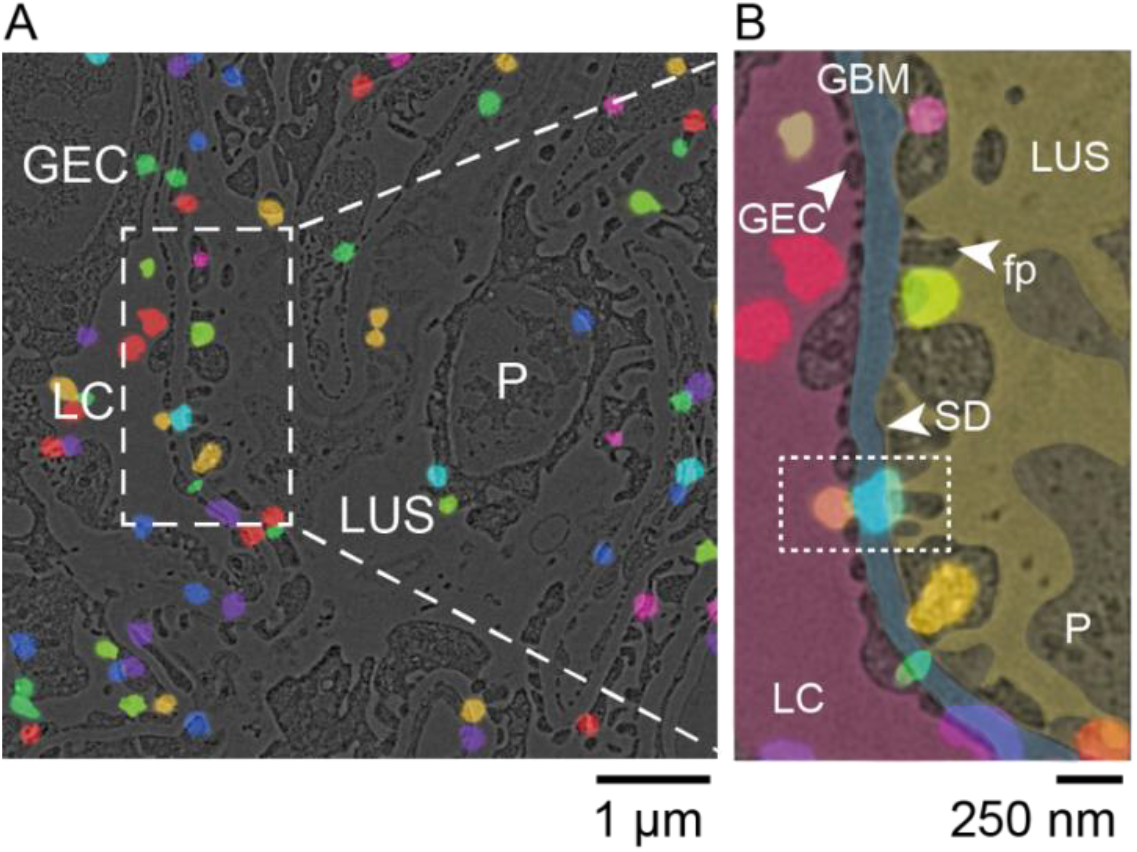
Super-resolution correlative light and electron microscopy displays the positions of albumin in the vicinity of the slit diaphragm. **(A)** Electron micrograph of a high-pressure frozen and freeze-substituted kidney section showing the luminal capillary (LC), glomerular endothelial cells (GEC), podocyte (P), and the lumen of the urinary space (LUS). The positions of fluorescent albumin clusters are shown in different colors. Albumin was labeled with Alexa Fluor 555. **(B)** Close-up of the micrograph shown in (a) with the manual segmentation of the areas of the LC (magenta) covered by GEC (light magenta), the glomerular basement membrane (GBM, light blue), and the podocyte with characteristic foot processes (fp and SD indicated by white arrowhead) in the LUS (yellow). The dotted line indicates an event of albumin passage through the filtration barrier.

Individual compartments of the glomerulus were analysed and the mean concentration of albumin clusters was calculated highest in the lumen of the capillaries (2.2 ± 1.0 clusters/µm^2^, n=28 field of views from five microbiopsies), lower in the GBM (1.6 ± 0.8 clusters/µm^2^, n=28 field of views from five microbiopsies), and lowest in the lumen of the urinary space (1.0 ± 0.9 clusters/µm^2^, n=28 field of views from five microbiopsies) (Fig. 4). One would anticipate that most of the albumin is held back by the filtration barrier, thus most of the albumin should be found in the LC and nearly nothing in the LUS. However, we still observe a substantial amount of albumin in LUS even though the SD seems intact in the electron micrograph. This can be partially attributed to the no-flow conditions (blood stream was interrupted) that the samples were subjected to before the experiment.

**Fig. 4:**
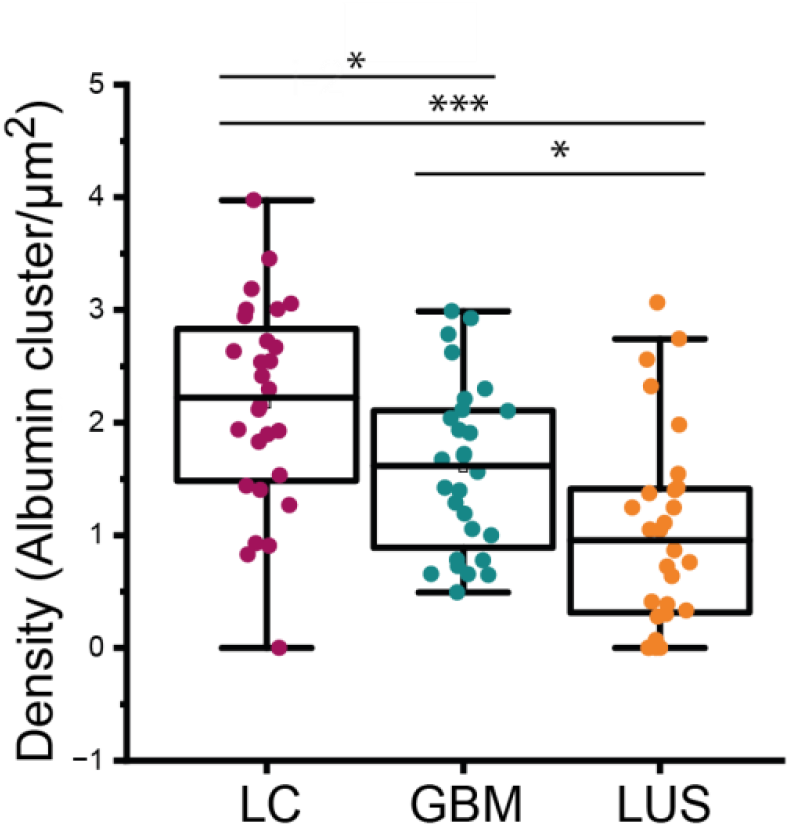
Statistical evaluation of the albumin density in the glomerular filtration barrier measured in albumin clusters per µm^2^. Statistical evaluation of the albumin density in the glomerular filtration barrier measured in albumin clusters per µm^2^ (for n=28). A linear reduction in the concentration can be observed. Paired t-test: LC v LUS p=5.3×10^−6^, GBM v LC p=0.0033, GBM v LUS p=4.56×10^−4^.

## Discussion

Visualizing and tracing the paths of molecules at nanometer resolution across the tissue with molecular resolution remains a challenging task. Here, we presented a feasibility study that demonstrates that the tracing of albumin is possible in the tissue context, i.e. through the glomerular filtration barrier. Although we only demonstrate this workflow for kidney tissue, it is transferable to other organs, particularly the brain. For example, it can be used to spot the leaking positions in the blood brain barrier or detect vessel tears associated with developing aneurysms. In addition, the workflow enables tracing the paths of viral particles or small molecules. Importantly, this workflow is not limited to cultured cells but is also applicable to volume electron microscopy. This study demonstrates that CLEM can effectively trace the paths of the fluorophores within tissue. Specifically, in tracing albumin within the kidney, we did not find traces of albumin in other subcellular compartments, such as within the podocytes, nor did we observe an increase vesicle transport or other physiological variations. This suggests that the SD is most likely responsible for the observed leakiness.

The current biophysical models of the renal filtration, the electrokinetic model postulates that under physiologic conditions albumin molecules never reach the SD [7,18,19]. Only under pathological conditions, such as podocyte diseases or conditions in which capillary flow ceases, do albumin molecules traverse the GBM and the SD. Our results also show a leakiness of the barrier at no-flow conditions showing that electrostatic forces are generated through the continuous flux, whereby negatively charged molecules are moved back to the capillaries by electrophoresis, which actively prevents the clogging of the glomerular filtration barrier. Thus, the streaming potentials impose a size and charge selectivity of the glomerular filtration, which prevent most of the filtrate from entering the filtration barrier at the endothelial cells [7]. Furthermore, the compression attributed from the podocytes on the GBM potentially reduces permeation speed. Despite those models the pathways by which proteins diffuse through the kidney filtration barrier remained unclear. In our analysis, we could not find albumin within the podocytes or other cell types. We find a linear reduction in the albumin concentration through the kidney filtration barrier. While the infiltration of albumin in the LUS may be attributed to the loss of the streaming potential it also shows that albumin can diffuse through the GBM and SD at no flow conditions. By leveraging the correlative approach of this study, which combines EM with high-resolution light microscopy, we can observe that the GBM is not uniformly penetrated by albumin clusters. Instead, albumin passage occurs at specific, localized areas where multiple albumin molecules cluster together to traverse the barrier. Unfortunately, the current resolution is insufficient to identify potential channels that might form between endothelial cells, extend through the GBM, and eventually open between the podocyte foot processes. This issue warrants further investigation, possibly through more targeted labelling of different serum proteins. However, this approach underlines its applicablity to studying protein distribution in other organs, such as the brain, where the exact mechanisms and locations of leakage remain unclear.

Cryo-electron tomography of vitreous lamella has emerged as the main technique to investigate the ultrastructure of cells [20]. However, only small regions of about 10 µm x (10-/)70 µm x 0.2 µm can be visualized, losing most of the surrounding tissue information due to the current preparation techniques. Volume electron microscopy offers a huge field of view and allow for a different quantification as shown here. CLEM is not trivial for conventional EM preparation methods as used in volume electron microscopy, because the harsh treatment. During the EM preparation, the proteins are denatured due to the acidic environment of the embedding material [21], which quench the GFP fluorescence. Previously it was shown that synthetic fluorophores do not degrade as severely and thus the heavy metal concentration can be increased to such an extent that the same sample can be used for TEM. Here we also show that this treatment is possible also for tissue and super resolution can also be achieved. Future experiments can be performed at flow conditions in kidney and methodology can be applied for any other tissue, where blood flow does not impact the proteins distributions.

## Data Availability Statement

## Acknowledgements and Funding

We thank the Frankfurt Center for Electron Microscopy and the Frankfurt Center for Advanced Light Microscopy for measurement time. We thank A. Habermann for assistance during plastic embedding. FG was supported by DFG (CRC 1192, GR3933/1-1). A.N.B., J.V.R. and M.H. were funded by Research Training Group iMOL (GRK 2566/1). A.S.F. was supported by the Deutsche Forschungsgemeinschaft (FR 1653/14-1 for U.H.E.).

## Author Contributions

### Author contributions

Conceptualization: MH, ASF

Formal analysis: ANB, JVR, UHE, ASF

Funding acquisition: FG, MH, ASF

Investigation: ANB, JVR, BF, FG, MH, ASF

Methodology: ANB, JVR, BF, UHE, AS, ASF

Project administration: ASF

Resources: ASF Supervision: FG, MH, ASF

Visualization: ANB, ASF

Writing – original draft: ANB, JVR, FG, MH, ASF

Writing – review & editing:

---

**Supplementary Figure 1:**
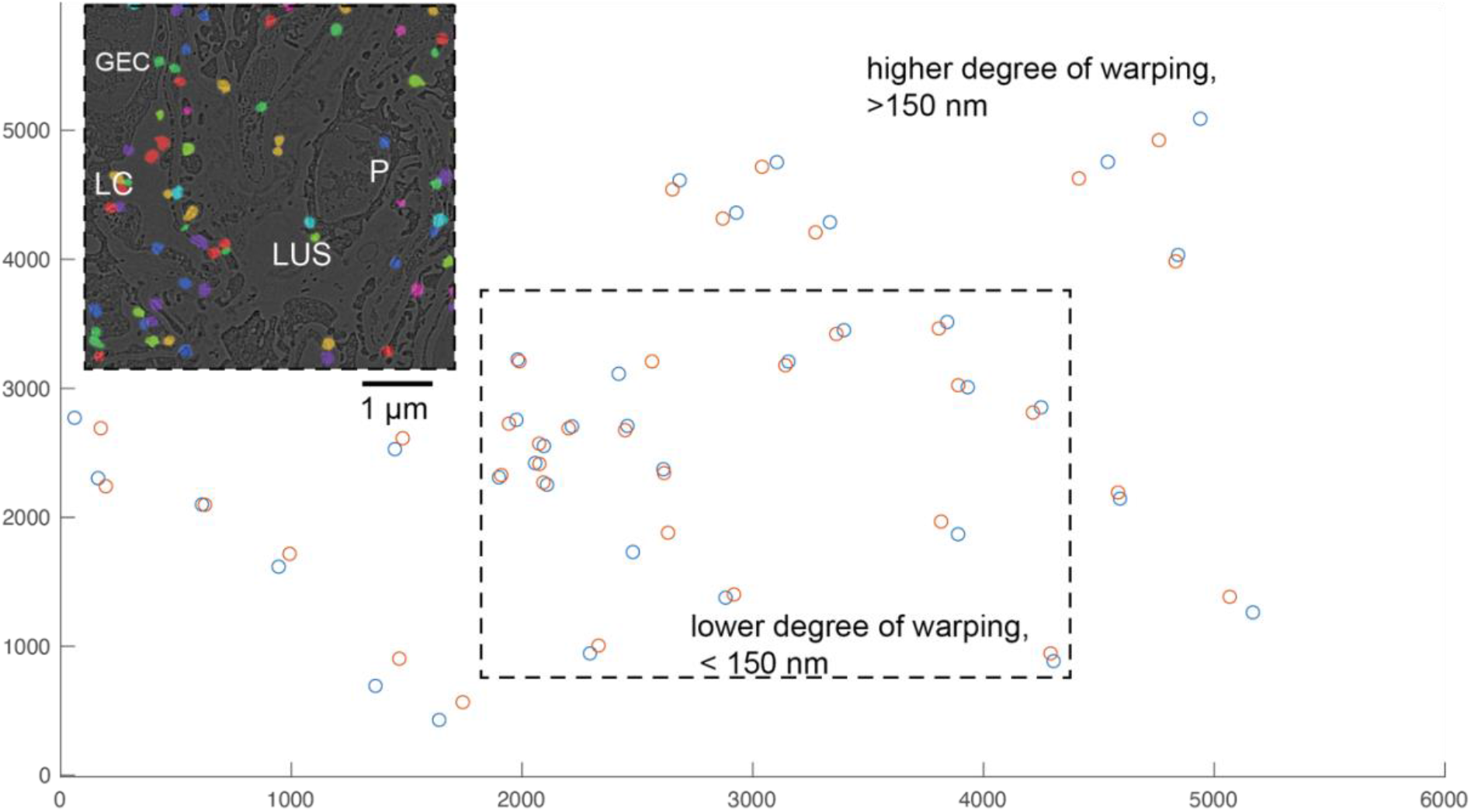
Example of omitted areas for the analysis. Accuracy of the image registration between super-resolution image (blue circles) and EM image (orange circles) was compared. Any areas of the sample that showed a higher degree of warping (>150 nm deviation) were omitted from the analysis, while regions that show less than ∼150 nm deviation were used (dashed boxed, inset for referent).

